# *Bacillus subtilis* reprograms the host transcriptome and rhizosphere microbiome with effects consistent with systemic responses to alkaline stress in garden pea

**DOI:** 10.1101/2025.09.13.676035

**Authors:** Ahmad H. Kabir, Asha Thapa, Md Rokibul Hasan, Mohammad G. Mostofa

## Abstract

Soil alkalinity severely limits legume growth, but the role of *Bacillus subtilis* in alkaline stress tolerance remains unclear in garden pea. We found that multiple garden pea genotypes inoculated with *B. subtilis* under alkaline stress showed host-specific improvements in growth parameters. Mechanistic analysis conducted on Sugar Snap showed improved nodulation, mineral status, and photosystem efficiency, while split-root assays showed responses consistent with systemic effects of *B. subtilis* in alkaline tolerance. Further, FeEDDHA partially reduced alkaline stress symptoms but did not fully restore nodulation. In contrast, *B. subtilis* increased rhizosphere Fe-chelating activity and improved nodulation, leading to stronger symbiotic recovery than inorganic Fe alone. This suggests that factors associated with *B. subtilis* inoculation, beyond Fe availability alone, may contribute to the observed recovery of nodulation. This is further supported by *in vitro* co-culture experiments showing enhanced growth of *R. leguminosarum* in the presence of *B. subtilis* under alkaline conditions, indicating potential microbial compatibility for coping with stress. RNA-seq analysis identified 958 upregulated and 1,134 downregulated genes in roots inoculated with *B. subtilis* under alkaline conditions. The upregulated genes were mostly involved in the sugar-mediated symbiotic association (*SWEET* and *GLUT*), pH homeostasis (*cation/H+ exchanger* and *ATPase*), and nutrient assimilation (*Ammonium transporter* and *Zn/Fe permease*). Microbial community analysis revealed that *B. subtilis* significantly altered bacterial alpha diversity under alkaline stress, whereas fungal alpha diversity remained unaffected. Further, *B. subtilis* reshaped the rhizosphere microbial community and enriched taxa such as *Pseudomonas*, *Pseudorhizobium*, and *Chaetomium*, which were potentially associated with responses to alkaline stress. Taken together, microbial interventions such as *B. subtilis* offer an effective strategy to boost legume tolerance to alkaline soils.

## INTRODUCTION

Soil alkalinity is a widespread abiotic stress that severely constrains crop productivity by limiting the solubility and availability of essential nutrients, particularly iron (Fe), manganese (Mn), and phosphorus (P) (Wang et al., 2025; Rengasamy, 2010). In alkaline soil, high pH disrupts root membrane transport systems, impairs chlorophyll (Chl) biosynthesis, and reduces photosynthetic efficiency, resulting in stunted growth and yield reduction (Thapa et al., 2015a). Legume crops are especially sensitive to alkaline stress due to their reliance on symbiotic nitrogen fixation, a process that is highly dependent on optimal nutrient uptake and balanced rhizosphere conditions (Tang et al., 2024). The increasing prevalence of alkaline soils in arid and semi-arid agricultural zones, exacerbated by irrigation with bicarbonate-rich water, underscores the urgent need for sustainable strategies to mitigate their impact (Gamalero et al., 2020).

To cope with soil alkalinity, plants employ a suite of adaptive mechanisms to maintain nutrient acquisition and cellular homeostasis. These include selective ion transport and proton pump activity to regulate the uptake of essential nutrients and cellular pH, respectively (Ku et al., 2022; Kabir et al., 2012). Another strategy involves the exudation of organic acids from roots, which acidify the rhizosphere and improve the solubility of otherwise inaccessible nutrients (Barrow & Hartemink, 2023). Morphological adaptations, such as the proliferation of deep and extensive root networks and the formation of microbial symbioses, further enhance nutrient uptake efficiency under alkaline conditions (Zhang et al., 2020; Chen et al., 2016).

Microbiome-based interventions represent a promising, environmentally sustainable strategy to strengthen plant resilience against alkaline stress. Beneficial microbes influence not only local root physiology but also systemic signaling networks that coordinate stress responses at the whole-plant level (Pantigoso et al., 2022; Ortíz-Castro et al., 2009). Such signals can reprogram the root transcriptome, activate defense mechanisms, and restructure microbial communities, making plants more adaptable to stressful conditions (Jamil et al., 2022; Pantigoso et al., 2022; Ruffel et al., 2008). Among plant growth-promoting rhizobacteria (PGPR), *Bacillus subtilis* has gained significant attention for its ability to withstand environmental stresses (Sharma et al., 2023), rapid colonization of roots, and production of diverse bioactive metabolites (Mahapatra et al., 2022; Hashem et al., 2019). It facilitates the secretion of organic acids to solubilize minerals, the production of siderophores to chelate Fe, the synthesis of phytohormones, and the modulation of antioxidant defenses (Hazarika et al., 2023; Chandwani et al., 2023). Despite these known benefits, little is understood about the roles of *B. subtilis* in regulating transcriptomes and root microbiomes in the context of soil alkalinity in legumes.

The garden pea (*Pisum sativum* L. subsp *sativum* var *sativum*), an important source of dietary micronutrients, is highly sensitive to alkaline stress (Thapa et al., 2025b; Dahl et al., 2012). Elevated soil pH can severely reduce yields by disrupting photosynthetic efficiency, which manifests as leaf chlorosis and growth penalty (Meisrimler et al., 2016). However, the potential of *B. subtilis* to alleviate alkaline stress in garden pea remains poorly understood. In this study, we investigated how *B. subtilis* inoculation modulates defense-signaling pathways, transcriptome reprogramming, and rhizosphere microbiome shifts to enhance alkaline stress tolerance in garden pea. Our findings provide novel insights into the potential of *B. subtilis*-based bioinoculants for improving legume productivity on alkaline soils, contributing to sustainable agriculture in marginal environments.

## MATERIALS AND METHODS

### Plant cultivation and growth conditions

Several garden pea cultivars, such as Sugar Snap, Oregon Sugar Pod, Sugar Ann, Little Marvel, Green Arrow, and Lincoln, were used in the initial screening of genotypes. Following the preliminary screening, Sugar Snap was selected for subsequent analyses because it exhibited a clear and consistent physiological response to both alkaline stress and *B. subtilis* inoculation, making it a suitable model for investigating the underlying mechanisms of alkaline stress tolerance. Seeds were surface-sterilized in 2% sodium hypochlorite for 5 min, rinsed three times with sterile water, and placed in trays for germination at 25 °C for 48 h. Uniform seedlings were transplanted into pots containing 500 g of a soil mixture (field soil from a pea-growing area combined with commercial potting mix at a 1:2 ratio). The treatment groups include four conditions using lime [7.5 g NaHCO□ and 4.5 g CaCO□ per pot] and *B. subtilis* (1 × 10^8^ CFU/mL; ARS Culture Collection–USDA, NRRL B-14596): (i) control (no lime, pH ∼6.5), (ii) alkaline (lime-amended soil, pH ∼7.8), (iii) alkaline+BS (lime-amended soil, pH ∼7.8, plus *B. subtilis* inoculum), and (iv) BS+ (no lime, pH ∼6.5, plus *B. subtilis* inoculum). For plant inoculation, 1 mL of the 1 × 10^8^ CFU mL□¹ inoculum was applied to the base of each seedling once at the time of transplantation. This NRRL B-14596 strain was selected based on a pilot study in which different *B. subtilis* strains were tested for their strong potential to enhance tolerance in pea plants exposed to alkaline stress. In addition to the NaHCO□ mixed with the soil from day 1, a 15 mM NaHCO□ solution was applied weekly throughout the 5-week growth period before data collection (Supplementary Fig. S1). For a targeted experiment, two additional treatments were included: FeEDDHA+ (lime-amended soil, pH ∼7.8, supplemented with FeEDDHA at 0.5 g/500 g soil) and FeEDDHA control (non-lime soil, pH ∼6.5, with FeEDDHA at 0.5 g/500 g soil). Soil pH was measured weekly by suspending 10 g of rhizosphere soil in 25 mL of deionized water (1:2.5, w/v), followed by thorough mixing and equilibration before measurement with a pH meter (Thermo Scientific Orion Star A214, USA). Plants were grown in a randomized complete block design under a 10 h light/14 h dark cycle (250 μmol m□² s□¹) at ∼25 °C in greenhouse conditions. Soil physicochemical properties are provided in Supplementary Table S1.

### Molecular detection of *B. subtilis* colonization in roots

DNA-based qPCR analysis was performed to assess *B. subtilis* colonization in Sugar Snap roots under different growth conditions. Root samples were rinsed twice in sterile phosphate-buffered saline, briefly vortexed, and washed twice with sterile water. Genomic DNA was extracted from approximately 0.2 g of root tissue using the cetyltrimethylammonium bromide (CTAB) method (Clarke, 2009). DNA quantity and purity were verified using a NanoDrop ND-1000 (Wilmington, USA), and all samples were normalized before qPCR. Amplification was conducted on a CFX96 Touch Real-Time PCR Detection System (Bio-Rad, USA) using *B. subtilis*-specific primers (forward: 5′-TCTGCTCGTGAACGGTGCT-3′; reverse: 5′-TTTCGCCTTATTTACTTGG-3′) (IARRPCAAS, 2013). Reactions were set up with iTaq™ Universal SYBR® Green Supermix (Bio-Rad, USA), with *P. sativum* GAPDH (Fw GTGGTCTCCACTGACTTTATTGGT, Rv TTCCTGCCTTGGCATCAAA) serving as the reference gene. Relative quantification of *B. subtilis* abundance was calculated using the 2□ΔΔCT method (Livak & Schmittgen, 2001). This assay was conducted using five biological replicates per treatment. Since the goal of this assay was to compare relative differences in *B. subtilis* colonization among treatments rather than to determine absolute bacterial numbers, quantification was performed using the 2□ΔΔCt method normalized to the pea housekeeping gene *GAPDH*. Plant GAPDH was used as an internal reference to normalize differences in host DNA input among root samples, allowing comparison of relative *B. subtilis* colonization across treatments.

### Measurement of morphological features and chlorophyll fluorescence kinetics

Shoot height and root length were measured for each plant using a measuring tape. Shoots and roots were then harvested and oven-dried at 70 °C for 3 days to determine dry weight. For nodule assessment, roots were gently washed to remove soil debris, and nodules were carefully detached, counted under a magnifying lens, and stored at −80 °C for subsequent analyses. Chlorophyll content was determined with a handheld SPAD meter (AMTAST, USA). Furthermore, chlorophyll fluorescence kinetics (OJIP), including the maximal photochemical efficiency of PSII (*F_v_*/*F_m_*) and the photosynthetic performance index (Pi_ABS), were recorded on the uppermost fully expanded leaves at three distinct points using a portable FluorPen FP 110 (Photon Systems Instruments, Czech Republic). For OJIP measurements, leaves were dark-adapted for 1 h before data acquisition. SPAD and OJIP measurements were taken once at the final harvest. All readings were collected between 10:00 am and 12:00 pm to minimize the effects of diurnal variation on photosynthetic parameters.

### Nutrient analysis in pea roots and leaves

Nutrient analysis was conducted on roots and young leaves. Root samples were excised, rinsed under running tap water, immersed in 0.1 mM CaSO_4_ for 10 min, and subsequently washed with deionized water. Young leaves were washed separately with deionized water. All samples were placed in labeled envelopes and oven-dried at 75 °C for 3 days. Elemental concentrations were determined using inductively coupled plasma mass spectrometry (ICP-MS) at the Agricultural and Environmental Services Laboratories, University of Georgia.

### CAS-based Fe-chelating activity assay

Fe-chelating activity in rhizosphere soil was determined using the chrome azurol S (CAS) assay following the method described by Himpsl and Mobley (2019). Briefly, soil samples were homogenized in 80% methanol and centrifuged at 10,000 rpm for 15 min. An aliquot of the resulting supernatant (500 μL) was mixed with an equal volume (500 μL) of CAS solution and incubated at room temperature for 5 min. Absorbance was measured at 630 nm, using 1 mL CAS reagent as the reference. Siderophore units were calculated as percentage siderophore units (%SU) using the formula: %SU = [(Ar − As) / Ar] × 100, where Ar represents the absorbance of the reference and As represents the absorbance of the sample. Since the CAS assay detects general Fe-chelating activity, the measured response was interpreted as siderophore-associated activity rather than identification or quantification of specific siderophore compounds.

### Determination of ammonium in roots

Ammonia concentration in root tissues was measured using Nessler’s reagent spectrophotometric method (Jeong et al., 2013). Fresh root tissue (100 mg) was homogenized in 5 mL of 2% (w/v) potassium chloride, and the homogenate was centrifuged at 12,000 rpm for 10 min. One milliliter of the resulting supernatant was mixed with 1 mL of Nessler’s reagent and incubated at room temperature for 15 min. Absorbance was recorded at 420 nm using a spectrophotometer. Ammonia content was calculated from a standard curve generated with known concentrations of ammonium sulfate and expressed on a fresh weight basis (µg g□¹FW).

### Split-root experiments

Plants were first grown for 2 weeks in sterile vermiculite before initiating the split-root assay in soil pots. Following transplant, split-root plants were cultivated for an additional 4 weeks before measurements were taken. The split-root setup was established using a single pot with a partitioned assembly, with modifications to the method described by Thilakarathna and Cope (2021). The taproot of each seedling was severed at the midpoint, and the lateral roots were evenly distributed between two compartments within the same pot, each filled with a 1:2 mixture of natural field soil and commercial potting mix. A solid plastic barrier was used to separate the compartments. The plastic barrier prevented soil and solution exchange between compartments. Each treatment consisted of five independent plants, with one plant representing one biological replicate. The plastic barrier extended through the soil profile to physically isolate the two root compartments and prevent root, soil, and solution exchange. The integrity of the barrier and physical separation of the root systems were visually verified at harvest. All plants were maintained under identical growth conditions, with soil alkalinity applied according to the treatment design. In each split-root system, one compartment received *B. subtilis* inoculation, while the other remained uninoculated.

### RNA-sequencing and bioinformatics analysis

RNA-seq analysis was performed on root tissues. Before RNA extraction, roots were rinsed twice with sterile water and vortexed for 10 s in sterile phosphate-buffered saline to remove surface debris. Cleaned roots were flash-frozen in liquid nitrogen and ground to a fine powder using a pre-chilled mortar and pestle. Total RNA was extracted using the SV Total RNA Isolation System (Promega Corporation, USA). Samples with RNA integrity number (RIN) values greater than 8 were selected, and 1 μg of RNA per sample was used for library preparation. Libraries were generated with the KAPA RNA HyperPrep Kit with Poly-A Selection (Kapa Biosystems, USA) and amplified using the KAPA HiFi HotStart Library Amplification Kit (Kapa Biosystems, USA). Sequencing was conducted on an Illumina NovaSeq 6000 platform (0.2 Shared S4, 150 bp paired-end) at the RTSF Genomics Core, Michigan State University. Overall, 90.0% of reads passed quality filtering with scores ≥ Q30. Per-sample read depth, distribution of transformed gene-expression, and sequencing quality metrics are provided in Supplementary Fig. S3.

Bioinformatics processing of raw FastQ files was performed using Partek Flow genomic analysis software (Partek, St. Louis, MO, USA). Adapter sequences and low-quality reads were removed with Trimmomatic (Bolger et al., 2014) to obtain high-quality clean reads. The filtered reads were aligned to the *Pisum sativum* reference genome (Ensembl Genomes 62) using HISAT2 (Kim et al., 2015). Read counts for each gene were quantified using HTSeq (Anders et al., 2015). Differential expression analysis was performed on raw gene-level count data using the Limma-Voom workflow. Raw counts were transformed to log2 counts per million (logCPM) with precision weights estimated by the voom function, followed by linear modeling and empirical Bayes moderation in limma. Genes with low expression (<10 counts across all samples) were excluded before differential expression analysis. Differentially expressed genes (DEGs) were identified using a false discovery rate (FDR) < 0.05 and an absolute log2 fold change ≥ 2. FPKM (per million mapped fragments) values were calculated for expression reporting and visualization but were not used as input for differential expression testing. Principal component analysis (PCA) was performed on normalized, log-transformed expression data to assess overall transcriptomic variation and sample clustering among treatments. Venn diagrams were generated using VennDetail (Hinder et al., 2017), and heatmaps were produced with the *pheatmap* package in R (Hu et al., 2021). Functional enrichment analysis of DEGs was conducted using ShinyGO 0.76.3 (Ge et al., 2020), with gene ontology (GO) annotations obtained from EnsemblPlants (Yates et al., 2022).

### Amplicon-based 16S and ITS sequencing in the rhizosphere

Rhizosphere microbial populations were characterized using Illumina amplicon sequencing, targeting the ITS region for fungi and the 16S rRNA gene for bacteria. Root samples were vortexed in sterile phosphate-buffered saline (PBS) for 10 s, rinsed twice with sterile water, and ∼0.2 g of cleaned tissue was used for DNA extraction with the CTAB method (Clarke, 2009). RNase and Proteinase K treatments were included to remove RNA and protein contaminants, respectively. Amplicon libraries were generated using primer pairs 341F (CCTACGGGNGGCWGCAG) and 805R (GACTACHVGGGTATCTAATCC) for the 16S rRNA gene, and ITS3 (GATGAAGAACGYAGYRAA) and ITS4 (TCCTCCGCTTATTGATATGC) for the ITS region. Sequencing was conducted on the Illumina NovaSeq 6000 platform (PE250). Raw reads were processed and trimmed with Cutadapt (Martin, 2011) and analyzed using the DADA2 pipeline (Callahan et al., 2016). Sequences identified as mitochondrial and chloroplast origin were removed. Taxonomic classification of amplicon sequence variants (ASVs) was performed using the UNITE database (Nilsson et al., 2019) for fungal ITS sequences and SILVA 138 (Quast et al., 2013) for bacterial 16S sequences. For sequencing quality control, raw reads were subjected to quality filtering, and sequencing quality was assessed based on effective read percentage, error rate, Q20 and Q30 scores, and GC content. The resulting clean reads showed 99.99–100% effectiveness, an error rate of 0.02%, Q20 values of 98.22–98.41%, and Q30 values of 94.56–94.92%. Microbial community structure was assessed by calculating Bray-Curtis distance matrices from Hellinger-transformed abundance data, followed by Principal Coordinate Analysis (PCoA) using the *phyloseq* and *vegan* packages in R. Diversity analyses included alpha and beta diversity metrics, relative abundance profiling, and non-parametric Kruskal–Wallis tests, with significance determined at P < 0.05 (McMurdie & Holmes, 2013).

### Microbial co-culture method

Interactions between *B. subtilis* and *R. leguminosarum* were evaluated on nutrient agar under control and alkaline conditions, with the latter supplemented with 15 mM NaHCO_3_ to adjust the pH to 7.8. *B. subtilis* NRRL B-14596 and *R. leguminosarum* viciae 3841 were first grown overnight in nutrient broth at 28 °C with shaking at 200 rpm. Cultures were then adjusted to 1 × 10□ CFU mL□¹, as determined by dilution plating and OD–CFU calibration curves established for each species. A 5 μL inoculum of each microbial strain was cultured in three configurations: *B. subtilis* alone, *R. leguminosarum* alone, and a mixed culture (1:1 ratio) of *B. subtilis* + *R. leguminosarum*. Plates were incubated for 7 days, after which colony growth was quantified using ImageJ software (National Institutes of Health, USA). The radius of each colony was measured at three independent points, and the mean value was calculated for subsequent analysis.

### Data analysis

Statistical analyses were performed in SPSS Statistics 20.0 (IBM Corp., Armonk, NY, USA) and R using the ggplot2 package for visualization. Prior to ANOVA, data were tested for normality (Shapiro–Wilk test) and homogeneity of variances (Levene’s test). When assumptions were violated, data were log- or square root-transformed; however, non-transformed values are shown when results remain unchanged. For experiments involving multiple pea genotypes and treatments, data were analyzed using a two-way ANOVA with genotype, treatment, and their interaction (genotype × treatment) as fixed factors. For analysis within a single genotype (Sugar Snap), one-way ANOVA was used with treatment as the independent factor. Following a significant overall ANOVA (P < 0.05), pairwise mean comparisons were performed using Fisher’s least significant difference (LSD) test at P < 0.05. In this study, the physiological assays were performed with five independent replications, whereas the RNA-seq and amplicon sequencing analyses were conducted with three independent replications.

## RESULTS

### Screening of genotypes for tolerance to alkaline stress with *B. subtilis*

Across genotypes, alkaline stress consistently reduced shoot height and SPAD scores compared to control plants, though the extent of recovery with *B. subtilis* varied (Table 1). In Sugar Snap and Oregon Sugar Pod, both shoot height and SPAD scores decreased significantly under alkaline stress but were similar to control levels when inoculated with *B. subtilis* with or without alkaline stress (Table 1). In Sugar Ann and Little Marvel, alkaline stress reduced both traits. Plants exposed to alkaline stress inoculated with *B. subtilis* improved shoot height, while BS+ produced the highest values in Little Marvel and Sugar Ann, exceeding the control (Table 1). In Little Marvel, SPAD scores improved under alkaline+BS and BS+ conditions, but BS+ matched control plants (Table 1). For the cultivar Sugar Ann, BS+ showed higher SPAD values than the control, but the difference was not statistically significant. In Green Arrow and Lincoln, alkaline stress significantly reduced shoot height and SPAD values relative to the control. Inoculation with *B. subtilis* under alkaline conditions partially alleviated these reductions, restoring both traits to levels comparable to the control. Similarly, plants inoculated with *B. subtilis* under non-alkaline conditions exhibited shoot height and SPAD values that were not significantly different from those of control plants (Table 1).

**Table 1.** Genotypic screening of garden pea capable of utilizing *B. subtilis* (BS) to cope with alkaline stress. Data represent means ± SDs of five independent biological samples. Different letters indicate statistically significant differences among treatments (*P* < 0.05).

| Genotypes | Shoot height (cm) |  |  |  | SPAD score |  |  |  |
| --- | --- | --- | --- | --- | --- | --- | --- | --- |
|  | Control | Alkaline | Alkaline+BS | BS+ | Control | Alkaline | Alkaline+BS | BS+ |
| Sugar Snap | 82±2.5a | 56±3.2b | 78±1.5a | 83±4.1a | 33±0.76a | 23±2.17b | 32±2.47a | 34±1.86a |
| Oregon Sugar Pod | 24±1.7a | 14±3.3b | 22±1.8a | 25±0.57a | 33±1.4a | 23±1.9b | 34±2.3a | 34±2.2a |
| Sugar Ann | 26±2.0ac | 21±0.57b | 24±0.57a | 32±3.6c | 39±1.9a | 33±2.7b | 36±1.7b | 36±2.4b |
| Little Marvel | 15±0.90a | 7±2.4b | 12±1.3a | 22±5.8c | 47±2.0a | 31±1.8b | 36±2.2c | 44±2.5a |
| Green Arrow | 24±1.1a | 19±1.9b | 20±1.7b | 28±3.0a | 47±1.7a | 39±1.9b | 42±1.5b | 48±1.3a |
| Lincoln | 19±1.0a | 15±0.57b | 14±2.0b | 19±1.5a | 34±0.36a | 30±1.5b | 31±1.5b | 37±3.8a |

### Effect of *B. subtilis* on physiological parameters and root colonization

Phenotypic observation showed that plants under alkaline had smaller shoots and roots compared to control (Fig. 1A). Plants inoculated with *B. subtilis* showed an improvement in shoot and root growth compared to alkaline stress, while plants inoculated with *B. subtilis* without stress appeared similar to control (Fig. 1A). The relative abundance of *B. subtilis* in roots was low in control and alkaline stress but significantly increased only in alkaline+BS conditions (Fig. 1B). Plant fresh weight was significantly reduced in alkaline compared to control, increased under alkaline stress with *B. subtilis* to values comparable to control, and remained similar to the plants solely inoculated with *B. subtilis* (Fig. 1C). Further, shoot height followed the same trend (Fig. 1D). Furthermore, root length (Fig. 1E) and root FW (Fig. 1F) were significantly reduced under alkaline stress compared to control. However, both parameters were restored to control levels under alkaline+BS conditions and remained comparable to control in BS+. In addition, root ammonium content was significantly affected by treatments. Control and BS+ plants showed the highest values, which were not significantly different from each other, while alkaline treatment had the lowest ammonium content (Fig. 1G). Plants inoculated with *B. subtilis* under alkaline stress partially restored ammonium levels, resulting in significantly higher values than alkaline alone but still lower than control and BS+ conditions (Fig. 1G). We also found that *F_v_*/*F_m_*and PI_ABS declined significantly in alkaline conditions compared to control, increased under alkaline stress inoculated with *B. subtilis* conditions to control levels, and remained unchanged in BS+ relative to control (Fig. 1H and 1I).

**Fig. 1.**
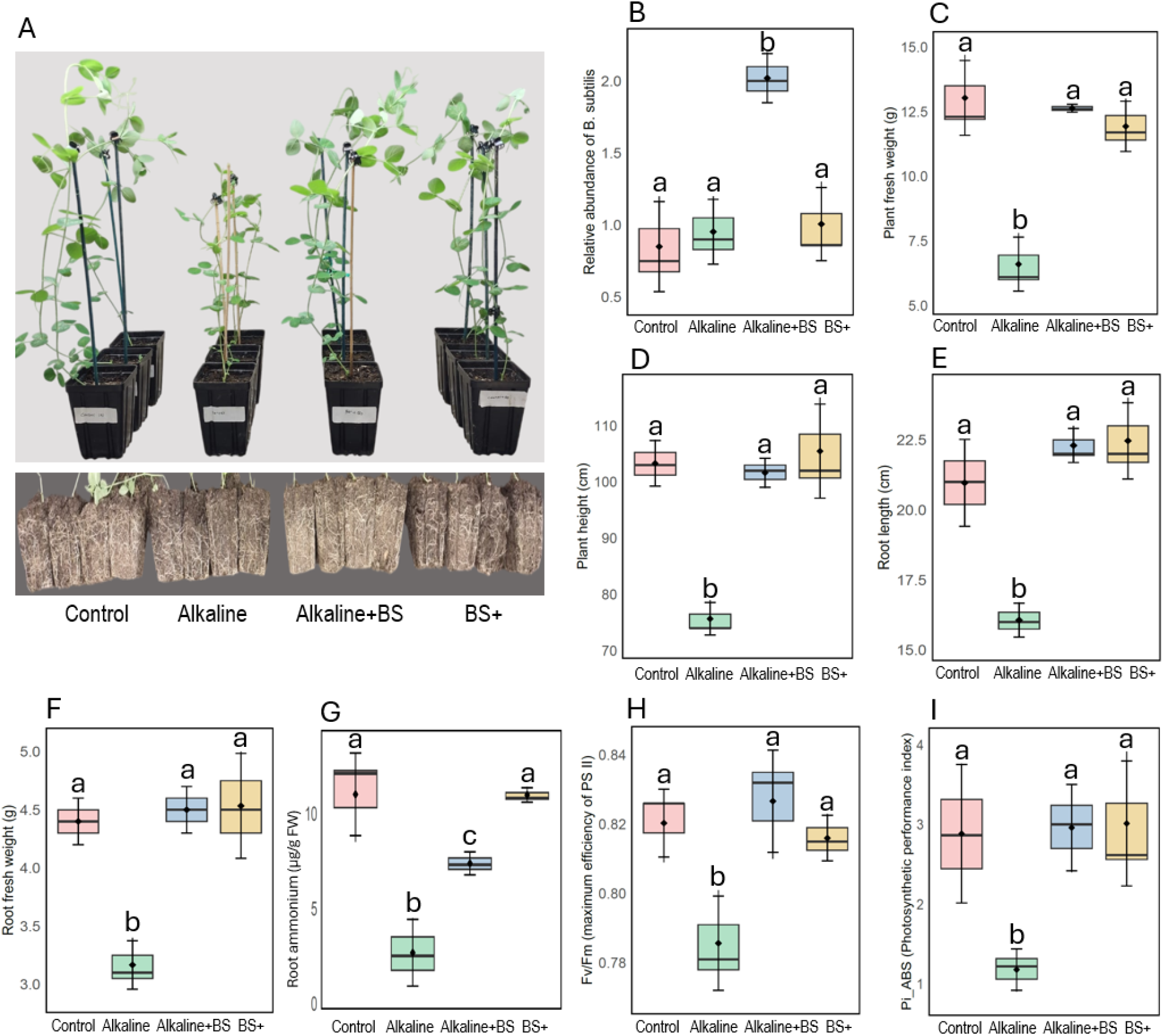
Effect of *Bacillus subtilis* on plant phenotypes (A), relative abundance of *B. subtilis* (BS) in the roots (B), plant fresh weight (C), plant height (D), root length (E), root fresh weight (F), root ammonium content (G), *F_v_*/*F_m_*: maximum efficiency of PSII (H), and PI_ABS: photosynthetic performance index (I) in Sugar Snap grown under four treatments: control, alkaline (pH 7.8), alkaline+BS, and BS+. Data represent means ± SD of five biological replicates (*n* = 5) on independent cultivations. Different letters above the box plots indicate statistically significant differences among treatments (*p* < 0.05, one-way ANOVA followed by Fisher’s LSD test).

### Effect of *B. subtilis* on nutrient concentrations in roots and leaves

ICP-MS analysis of nutrient contents showed a variation in their concentrations in the root and leaf tissues of garden peas grown with or without *B. subtilis* under alkaline conditions (Table 2). In alkaline-stressed plants, there was a decrease in concentrations of Fe, Mn, and N in both root and leaf tissues. However, the addition of *B. subtilis* to alkaline-stressed plants recovered these nutrient contents in both root and leaf tissues and was similar to control plants. Additionally, *B. subtilis* inoculated with control plants showed Fe, Mn, and N concentrations in the roots and leaves of garden peas that were similar to those in the control and alkaline + BS treatments. However, there were no significant changes in the amount of S in the root tissues under alkaline conditions compared to the control. There was a decrease in S content in leaf tissues under such conditions, and it was recovered when *B. subtilis* was added (Table 2). Similarly, there was a significant decrease in magnesium (Mg) content in roots of alkaline-stressed plants; however, there was no change in Mg content in leaf tissues under all four treatment conditions. Additionally, under alkaline conditions, calcium (Ca) concentrations in both the roots and leaves of garden peas decreased significantly, while inoculation of alkaline-stressed plants with *B. subtilis* increased Ca content in both tissues (Table 2).

**Table 2.** Changes in nutrient concentrations in roots and leaves of pea plants inoculated with or without *B. subtilis* (BS) under control and alkaline soil conditions. Data represent means ± SDs of five independent biological samples. Different letters indicate statistically significant differences among treatments (*P* < 0.05).

| Elements | Roots |  |  |  | Leaves |  |  |  |
| --- | --- | --- | --- | --- | --- | --- | --- | --- |
|  | Control | Alkaline | Alkaline+BS | BS+ | Control | Alkaline | Alkaline+BS | BS+ |
| Fe (ppm) | 1011±72.8a | 660±96.8b | 997±130.4a | 999±42.1a | 202±33.1a | 137±22.0b | 205±17.6a | 196±23.2a |
| Mn (ppm) | 70±5.5a | 39±11.6b | 100±25.0a | 114±27.3a | 49±7.1a | 31±3.6b | 37±5.8a | 46±10.6a |
| S (%) | 0.10±0.01a | 0.09±0.01a | 0.10±0.005a | 0.09±0.01a | 0.22±0.3a | 0.13±0.01b | 0.23±0.02a | 0.24±0.01a |
| Mg (%) | 0.29±0.05a | 0.16±0.01b | 0.36±0.07a | 0.28±0.05a | 0.15±0.2a | 0.09±0.01a | 0.14±0.02a | 0.16±0.03a |
| Ca (%) | 0.76±0.08a | 0.51±0.04b | 1.01±0.03a | 0.99±0.09a | 0.28±0.03a | 0.18±0.02b | 0.49±0.06c | 0.53±0.11c |
| N (%) | 3.2±0.26a | 2.4±0.18b | 3.2±0.22a | 3.4±0.57a | 7.0±1.19a | 5.5±1.33b | 7.1±0.68a | 7.6±1.04a |

### Comparative performance of *B. subtilis* and inorganic Fe in alkaline soil

The effects of *B. subtilis* inoculation and inorganic Fe supplementation on pea growth under alkaline conditions were investigated and compared (Fig. 2A). The relative abundance of *B. subtilis* in roots was low in control, alkaline, alkaline+Fe, and inorganic Fe plants (Fig. 2B). In contrast, alkaline+BS conditions showed a significant increase in *B. subtilis* abundance as compared to other treatments without *B. subtilis* additions (Fig. 2B). Rhizosphere Fe-chelating activity measured by the CAS assay differed markedly among treatments (Fig. 2C). Before applying the inoculum to plants, we first tested whether *B. subtilis* maintained consistent Fe-chelating activity under control and alkaline nutrient conditions. Fe-chelating activity by the *B. subtilis* inoculum showed no statistically significant differences between treatments (Supplementary Fig. S2). Alkaline conditions caused a significant decline in Fe-chelating activity in the rhizosphere compared to controls. When *B. subtilis* was introduced under alkaline conditions, Fe-chelating activity increased significantly compared to alkaline conditions, reaching levels statistically comparable to the control (Fig. 2C). Interestingly, supplementation with inorganic Fe under alkaline conditions resulted in no significant increase compared to alkaline stress, which remained significantly lower than *B. subtilis*-treated plants (Fig. 2C). Inorganic Fe under alkaline conditions did not restore Fe-chelating activity to control levels, whereas alkaline+BS fully recovered Fe-chelating activity (Fig. 2C).

**Fig. 2.**
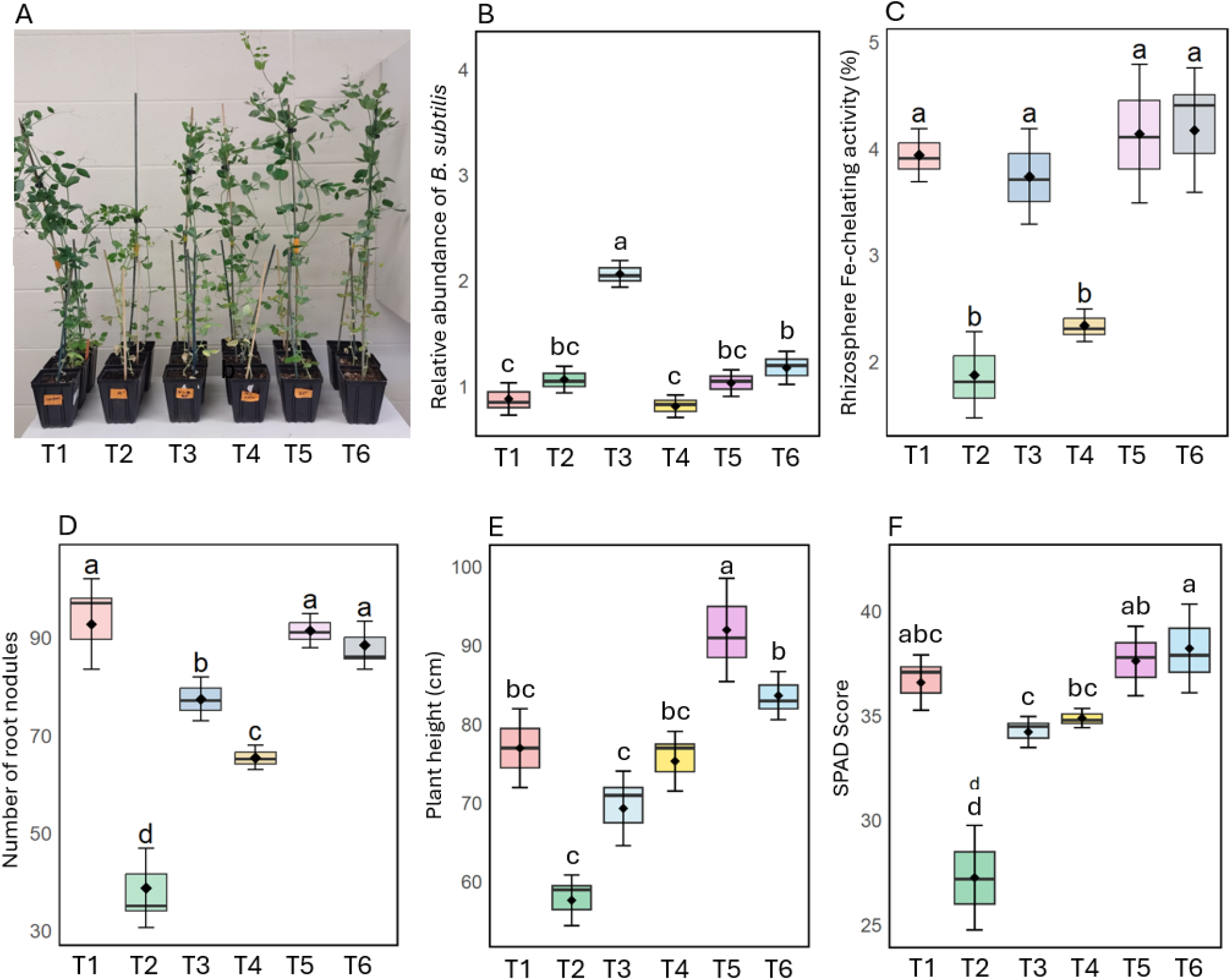
Growth performance and physiological responses of pea plants under different treatments (T1: control, T2: alkaline (pH 7.8), T3: alkaline + *B. subtilis* BS, T4: alkaline + inorganic Fe, T5: BS+, and T6: inorganic Fe) on phenotype of plants from each treatment (A), the relative abundance of *B. subtilis* in the roots (B), rhizosphere Fe-chelating activity (C), number of root nodules (D), plant height (E), and leaf SPAD score (F). Data represent means ± SD of five biological replicates (*n* = 5). Different letters above the box plots indicate statistically significant differences among treatments (*p* < 0.05, one-way ANOVA followed by Fisher’s LSD test).

Alkaline stress had a significant negative effect on nodule formation in pea roots compared to the control (Fig. 2D). Inoculation with *B. subtilis* under alkaline conditions partially rescued nodulation, resulting in significantly higher nodule numbers than in alkaline-stressed plants without inoculation. In contrast, supplementation with inorganic Fe under alkaline conditions provided only a modest improvement, and the increase remained lower than that observed in *B. subtilis*–inoculated alkaline plants (Fig. 2D). Furthermore, plants inoculated with *B. subtilis* under non-alkaline conditions or supplemented with inorganic Fe exhibited nodule numbers comparable to the control (Fig. 2D). Under alkaline stress, plant height decreased significantly compared to the control (Fig. 2E). Inoculation with BS or supplementation with inorganic Fe under alkaline stress significantly increased shoot height relative to alkaline conditions, with values approaching those of the control. Plants inoculated with *B. subtilis* exhibited plant height similar to controls, whereas inorganic Fe supplementation increased plant height slightly above control values (Fig. 2E). Furthermore, SPAD scores under alkaline stress were significantly reduced in relation to those in control plants (Fig. 2F). Both alkaline+BS and alkaline+Fe treatments restored SPAD scores to levels significantly higher than alkaline-stressed plants. Plants solely inoculated with *B. subtilis* or inorganic Fe showed similar SPAD scores to those of control plants (Fig. 2F).

### Split-root assay for identifying local/systemic response

A split-root system was used to distinguish between local and systemic effects of *B. subtilis* inoculation underlying alkaline stress tolerance in pea plants (Fig. 3A–F). In split-root experiments, root length and root fresh weight did not differ significantly between compartments across all treatments (Fig. 3A–B). However, plants exposed to alkaline stress in both compartments (SR2) showed no significant reduction in root length and root fresh weight compared with the control treatment (SR1) (Fig. 3A-B). In contrast, SR3 (alkaline/alkaline+BS) and SR4 (alkaline+BS/alkaline+BS) plants showed substantial improvements in root length and root fresh weight relative to SR2 (alkaline/alkaline) plants (Fig. 3A–B). Further, shoot traits responded strongly to treatment combinations (Fig. 3C–D). Among all treatments, SR2 consistently exhibited the lowest shoot growth responses compared to SR1. Plants exposed to *B. subtilis* in both compartments (SR4) or in one compartment under alkaline stress (SR3) showed significantly greater shoot height and shoot fresh weight than plants under alkaline stress without inoculation (Fig. 3C–D). Furthermore, leaf SPAD and Fv/Fm were significantly reduced under alkaline stress without inoculation (SR2), whereas plants inoculated with *B. subtilis* in one or two compartments with or without alkaline stress (SR3–SR6) showed a significant increase in SPAD and Fv/Fm levels compared to SR2 plants (Fig. 3E-F). Under non-alkaline conditions, SR5 and SR6 showed similar shoot height, shoot fresh weight, SPAD score, and Fv/Fm, with no significant differences between these treatments (Fig. 3C–F).

**Fig. 3.**
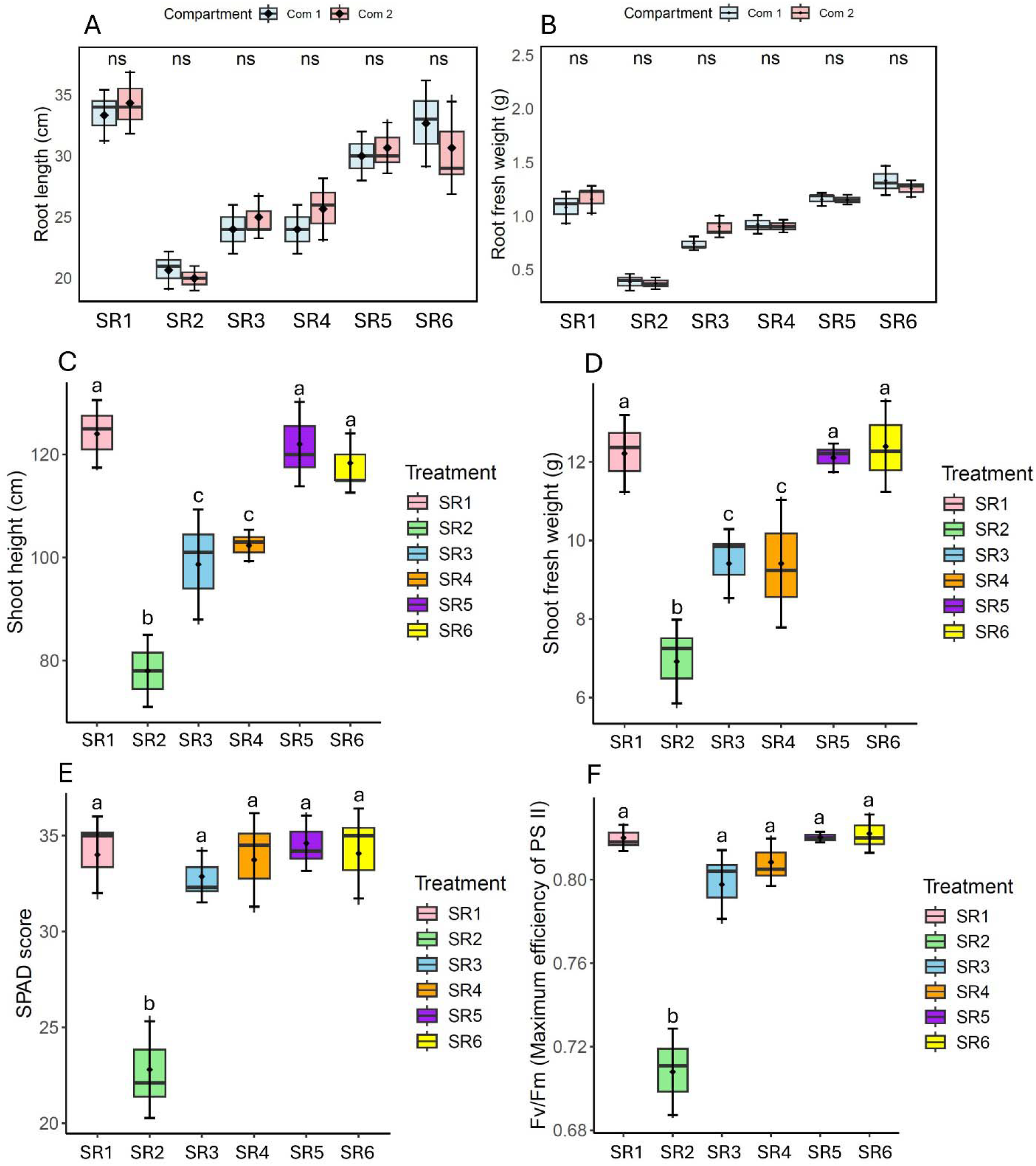
Effects of different split-root treatments (SR1: control/control, SR2: alkaline/alkaline, SR3: alkaline/alkaline+BS, SR4: alkaline+BS/alkaline+BS, SR5: control/control+BS, SR6: control+BS/control+BS) on root length (A), root fresh weight (B), shoot height (C), shoot fresh weight (D), SPAD score (E), and *F_v_*/*F_m_*: maximum efficiency of PSII (F) of pea plants cultivated in two compartments (com 1 and com 2). In treatments involving localized bacterial inoculation, one compartment received the *B. subtilis* inoculum while the other remained uninoculated; in bilateral inoculation treatments, both compartments received the inoculum. Data are means ± SD from five biological replicates (*n* = 5), with letters indicating statistical differences based on one-way ANOVA followed by Fisher’s LSD test (*p* < 0.05).

### Interaction between *B. subtilis* and *R. leguminosarum* in co-culture

The effect of co-culturing *B. subtilis* with *R. leguminosarum* was assessed under both control and alkaline conditions over three, five, and seven days (Fig. 4A-4B). Under non-alkaline conditions, colony sizes of *R. leguminosarum* and *B. subtilis* increased progressively over time in both single and co-culture treatments. On day three, no significant differences in colony size were observed between single and co-culture for either species. By day five and day seven, both species exhibited larger colony diameters compared to day three, but differences between single and co-culture remained statistically non-significant (Fig. 4A). Under alkaline conditions, colony sizes of both *R. leguminosarum* and *B. subtilis* were consistently smaller compared to control pH across all time points. On day three, growth reduction was evident in both species relative to their respective control pH counterparts, and coculture significantly altered the colony size of *R. leguminosarum* with an increase in colony area (Fig. 4B). Similar patterns were observed on days five and seven, with gradual increases in the colony diameter of *R. leguminosarum* over time, and significant differences between the single- and co-culture treatments under alkaline stress. However, there were no changes in colony growth of *B. subtilis* over three, five, and seven growing days between single and co-culture conditions (Fig. 4B). Overall, colony expansion for both species was slower in alkaline conditions than in control pH, and co-culturing confers a measurable growth advantage for only *R. leguminosarum* under alkaline conditions (Fig. 4B).

**Fig. 4.**
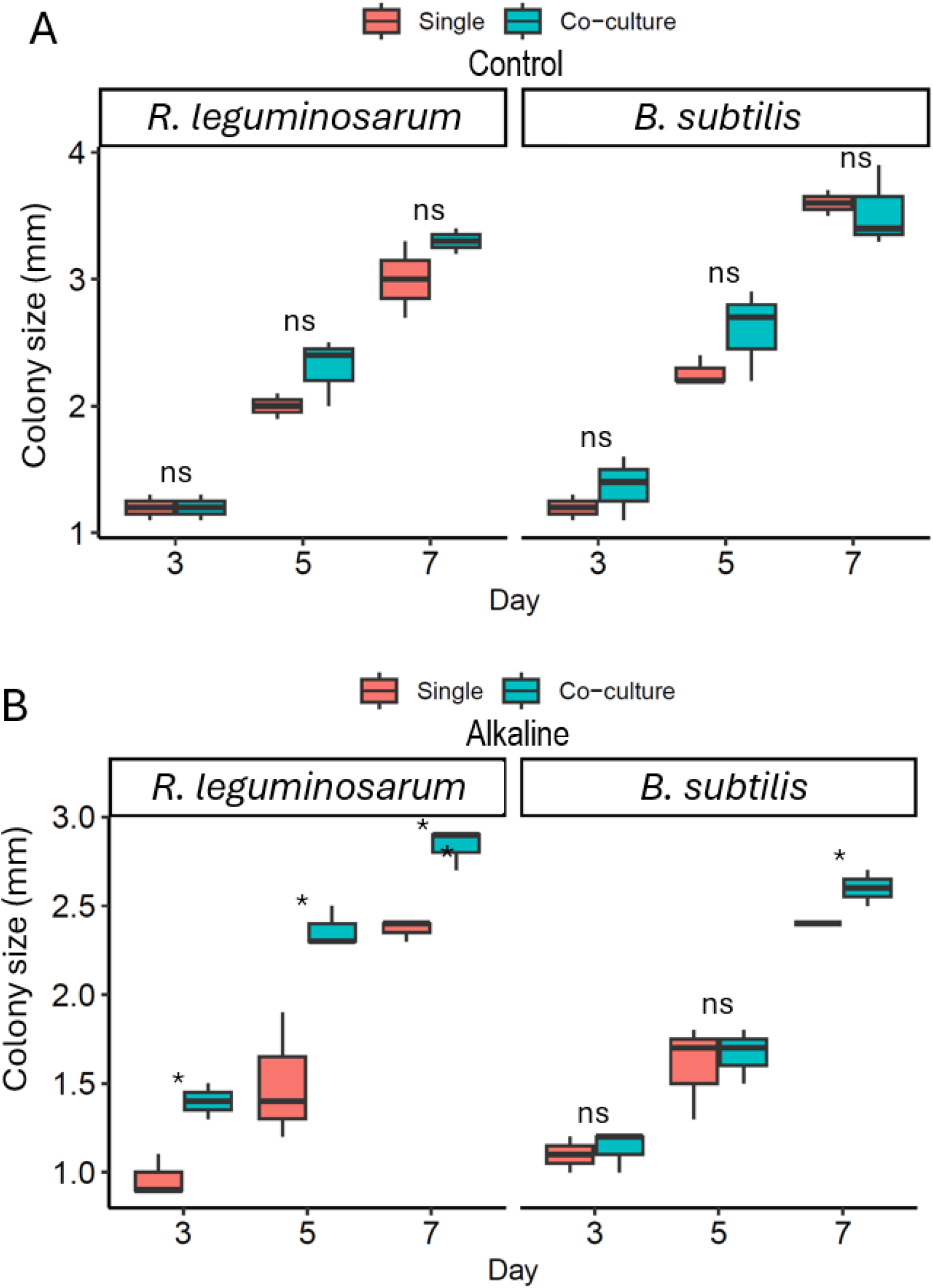
Effect of single and co-culture growth on colony size of *R. leguminosarum* and *B. subtili* under control (A) and alkaline conditions, pH 7.8 (B) over 3, 5, and 7 days. Data are means ± SD from five biological replicates (*n* = 5), with significance (* = *P* < 0.05, ns = not significant) determined by *t*-test comparison between single and co-culture within each time point.

### Transcriptomic changes in roots

Principal component analysis (PCA) revealed four distinct clusters, indicating that each treatment generated a unique transcriptomic profile (Fig. 5A). In Venn diagram analysis, alkaline+BS treatment had a unique set of upregulated (958) and downregulated (1134) genes in the roots compared to alkaline stress (Fig. 5B-5C). Specifically, alkaline+BS exhibited a large number of uniquely upregulated genes not shared with alkaline or BS+, along with a smaller subset of genes overlapping with control-related profiles (Fig. 5A). Heatmap visualization (Fig. 5D–G) highlighted key functional categories of differentially expressed genes (DEGs). In the alkaline+BS treatment, numerous genes related to symbiotic association (e.g., sugar phosphate transporter, glucose transporter GLUT, Sugar transporter SWEET, MF sugar transporter–like, Sugar transport protein STP, UDP–sugar pyrophosphorylase) were significantly upregulated compared to alkaline stress (Fig. 5D; Supplementary Table S2). Genes associated with nutrient transport and assimilation (e.g., ammonium transporter, nitrite and sulphite reductase, inorganic pyrophosphatase, amino acid transporter, Cys/Met metabolism, Iron/zinc purple acid phosphatase, malic oxidoreductase, SulP transporter, Zinc/iron permease, MFS transporter) were also more highly expressed in alkaline+BS compared to alkaline stress (Fig. 5E). Under alkaline conditions without *B. subtilis*, these genes generally exhibited reduced expression compared to control plants (Fig. 5E). Furthermore, redox and osmotic adjustment genes (e.g., glutaredoxin, oxidative stress regulator, glutathione synthase, thioredoxin, glutathione *S*-transferase, ferritin-like superfamily, dehydrin, aquaporin transporter) were induced in alkaline+BS relative to the plant solely stressed with alkaline (Fig. 5F). Lastly, several genes (*isoprenoid synthase*, *ethylene-responsive TF*, *cation/H+ exchanger*, *ABC-2 type transporter*, *ABC transporter type 1*, *root meristem growth factor 3*, *AP2/ERF ethylene responsive factor*, *oxoglutarate*, *small auxin-up RNA*, *ATPase*) related to growth and stress response showed significant upregulation due to B. subtilis under alkaline stress related to alkaline soil (Fig. 5G; Supplementary Table S2).

**Fig. 5.**
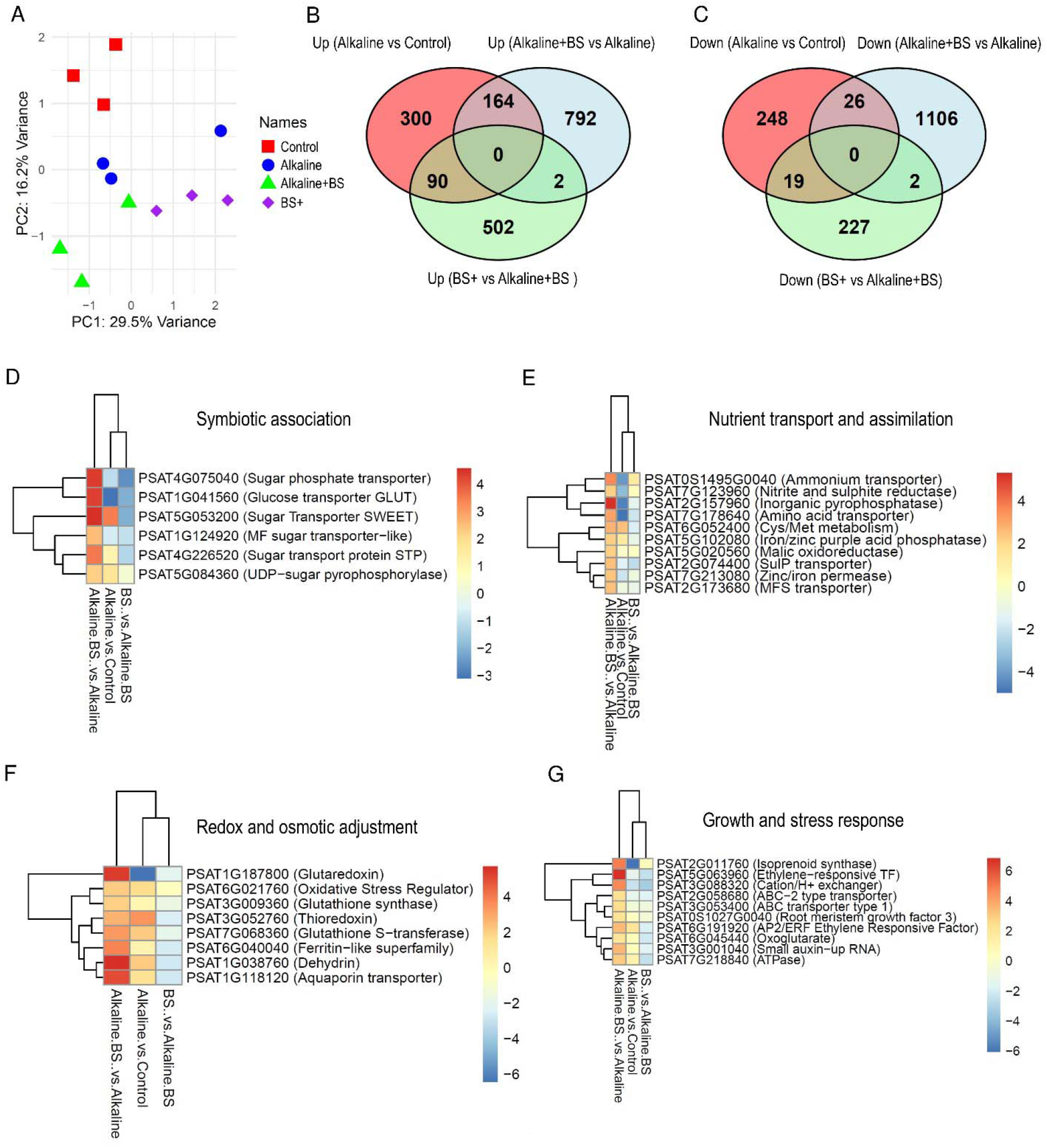
Transcriptomic changes in pea roots under control, alkaline, alkaline + *B. subtilis* (alkaline+BS), and BS+ treatments. (A) Principal component analysis (PCA) plot showing separation of transcriptome profiles among treatments, (B) Venn diagram of upregulated genes, (C) Venn diagram of downregulated genes, (D–G) Heatmaps showing selected differentially expressed genes (DEGs) grouped by functional categories. Color scale represents normalized expression values (log fold change), with red and blue indicating upregulation and downregulation, respectively. Data are based on three independent biological replicates per treatment (*n* = 3).

### Gene ontology enrichment analysis

Gene ontology (GO) enrichment analysis of the alkaline+BS versus alkaline comparison revealed distinct functional categories associated with upregulated and downregulated genes (Fig. 6A-B). Among the upregulated genes, the most significantly enriched GO terms were related to responses to water, responses to acid chemicals, and responses to inorganic substances, indicating activation of stress-responsive pathways under *B. subtilis* treatment. Additional enriched terms included oxidoreductase activity, protein processing in the endoplasmic reticulum, and several membrane- and transporter-associated functions, suggesting enhanced cellular transport and redox regulation (Fig. 6A). Conversely, downregulated genes were strongly enriched for GO categories associated with oxygen binding, heme and tetrapyrrole binding, ion binding, and defense responses, indicating repression of oxidative and defense-related processes in the alkaline+BS treatment. Several broad metabolic and regulatory categories, including regulation of cellular processes, cellular metabolic processes, nitrogen (N) compound metabolism, and macromolecule biosynthesis, were also significantly enriched among the downregulated genes, reflecting a reorganization of metabolic priorities (Fig. 6B).

**Fig. 6.**
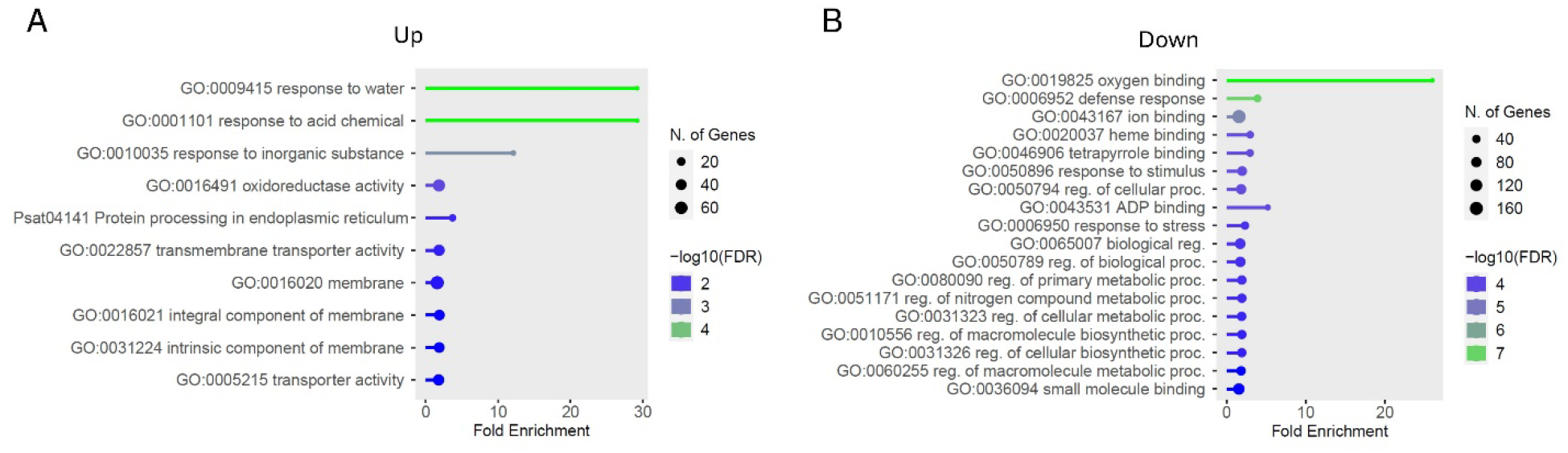
Gene ontology (GO) enrichment analysis of differentially expressed genes in the alkaline+BS *vs.* alkaline comparison using ShinyGO. (A) Top enriched GO terms for upregulated genes and (B) Top enriched GO terms for downregulated genes associated with biological process. Bubble size represents the number of genes in each GO term, color indicates log10(FDR), and the x-axis shows fold enrichment. Data are based on three biological replicates per treatment (*n* = 3).

### Rhizosphere microbial community structure

Principal coordinate analysis (PCoA) based on Bray–Curtis dissimilarity indicated that plants grown under non-stressed conditions, with or without *B. subtilis*, occupied a similar region of ordination space (Fig. 7A). In contrast, alkaline and alkaline+BS samples were clearly separated from the non-stressed groups and from each other (Fig. 7A). Alpha diversity analysis showed modest variation across treatments, but overall differences were limited. Particularly, alkaline stress altered bacterial richness, while B. subtilis modestly shifted richness patterns; however, these trends were not strongly treatment-specific (Fig. 7B). In contrast, Simpson diversity did not differ significantly among treatments (Fig. 7B). At the family level, *Pseudomonadaceae*, *Gemmatiomonadaceae*, and *Flavobacteriaceae* showed statistically significant differences in relative abundance among the treatments (Fig. 7C). *Pseudomonadaceae* and *Flavobacteriaceae* were markedly enriched in the rhizosphere of plants inoculated with *B. subtilis* under alkaline stress compared to alkaline stress only. Other dominant families such as *Gemmatiomonadaceae*, *Rhizobiaceae*, and *Sphingomonadaceae* exhibited treatment-dependent shifts but without statistical significance (Fig. 7C). At the genus level, *Variovorax*, *Shinella*, *Pseudorhizobium*, *Pseudomonas*, *Novosphingobium*, *Flavobacterium*, *Brevundimonas*, and *Bosea* displayed statistically significant variation across treatments (Fig. 7D). Particularly, *Pseudomonas* and *Pseudorhizobium* showed a significant enrichment in alkaline-exposed conditions inoculated with *B. subtilis* relative to the conditions solely exposed to alkaline stress. In contrast, *Shinella* was enriched under alkaline stress, while *Bosea* and *Variovorax* were predominant in control plants (Fig. 7D).

**Fig. 7.**
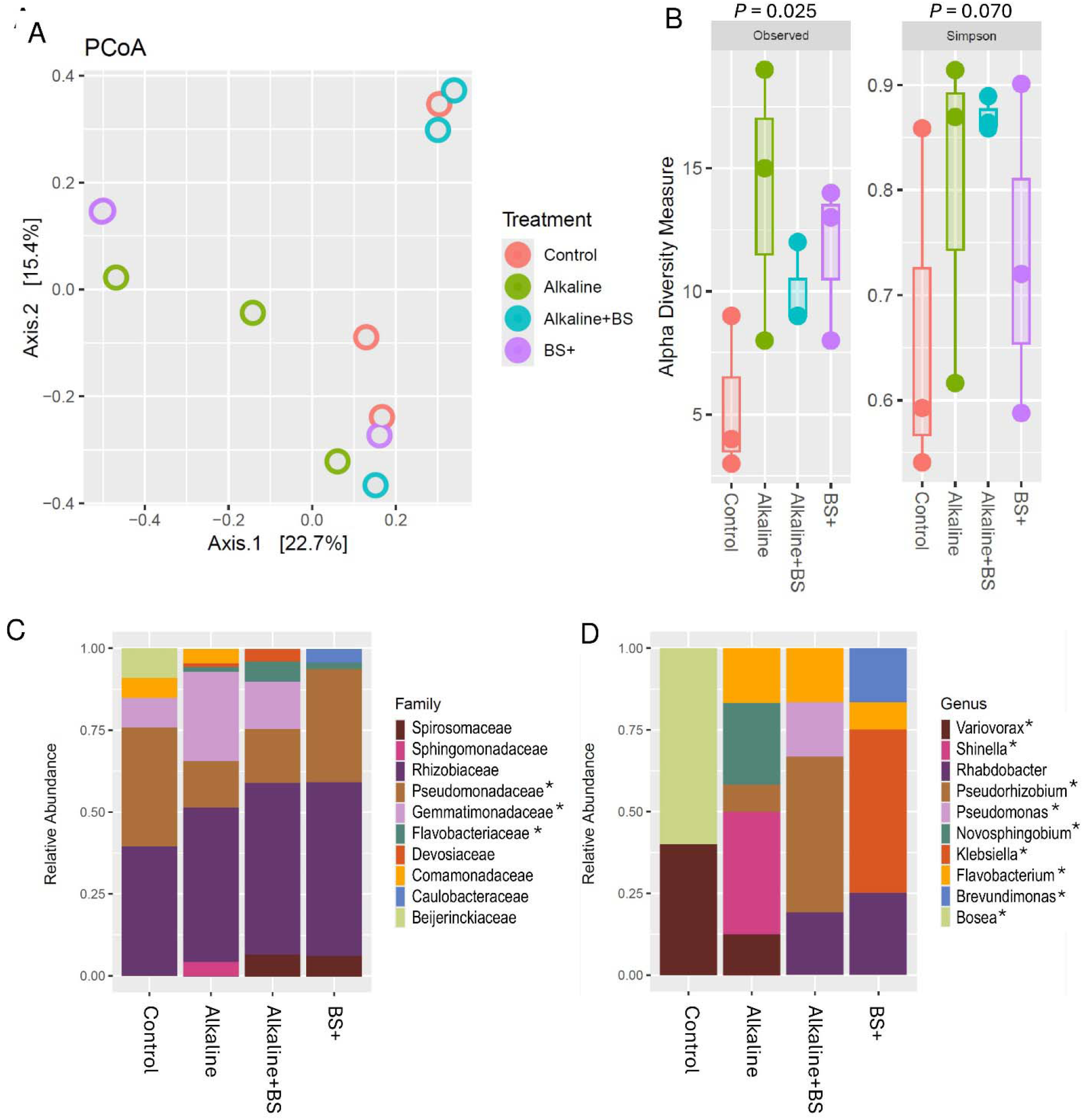
Rhizosphere bacterial community structure in Sugar Snap under different treatments: control, alkaline, alkaline+*B. subtilis* (BS), and BS+. Data represent (A) principal coordinate analysis (PCoA) plot based on Bray–Curtis dissimilarity showing separation of bacterial communities among treatments, (B) alpha diversity indices (Observed richness and Simpson index; box plots show median, interquartile range, and individual replicates, (C) relative abundance of dominant bacterial families, and (D) relative abundance of dominant bacterial genera. Asterisks (*) indicate taxa that showed statistically significant differences among treatments based on Kruskal-Wallis (*P* < 0.05). Data are from three biological replicates per treatment (*n* = 3).

In this study, PCoA of ITS showed distinct clustering of rhizosphere fungal communities among the treatment groups (Fig. 8A). Particularly, control and BS+ grouped closely, indicating minimal differences in fungal composition under non-stress conditions. In contrast, alkaline and alkaline+BS occupied separate positions from control, with alkaline+BS partially shifting toward the control/BS+ cluster, indicating that *B. subtilis* inoculation under alkaline stress altered fungal community structure (Fig. 8A). In our analysis, observed richness and Simpson diversity index showed no significant changes due to B. subtilis in the presence or absence of soil alkalinity (Fig. 8B). Among the families, *Halosphaeriaceae* and *Cladosporiaceae* exhibited lower relative abundance, whereas *Chaetomiaceae* showed higher abundance under alkaline stress inoculated with *B. subtilis* compared to alkaline stress (Fig. 8C). In contrast, the dominant family *Aspergillaceae* displayed broadly similar abundance between the two treatments. Among significant genera, *Pseudallescheria* and *Chaetomium* showed significant enrichment in the rhizosphere of plants inoculated with *B. subtilis* under alkaline stress compared to alkaline stress (Fig. 8D).

**Fig. 8.**
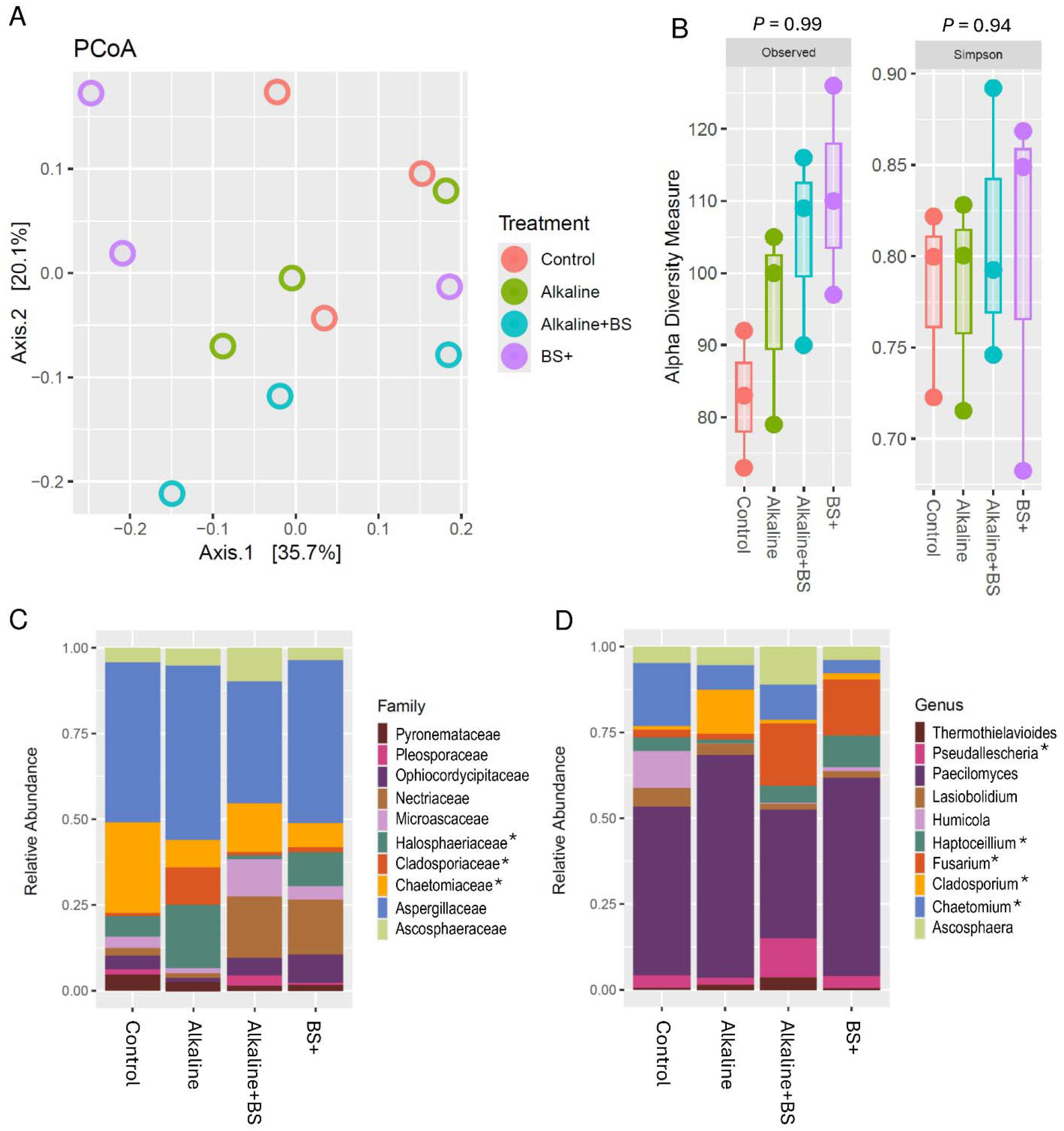
Rhizosphere fungal community structure in Sugar Snap under different treatments: control, alkaline, alkaline+*B. subtilis* (BS), and BS+. Data represent (A) principal coordinate analysis (PCoA) plot based on Bray–Curtis dissimilarity showing separation of fungal communities among treatments, (B) alpha diversity indices (Observed richness and Simpson index); box plots show median, interquartile range, and individual replicates, (C) relative abundance of dominant fungal families, and (D) relative abundance of dominant fungal genera. Asterisks (*) indicate taxa that showed statistically significant differences among treatments based on Kruskal-Wallis (*P* < 0.05). Data are from three biological replicates per treatment (*n* = 3).

## DISCUSSION

Alkaline stress in soil is a major constraint to legume productivity, primarily due to reduced availability of essential micronutrients (Gong et al., 2020). In this study, alkaline stress significantly disrupted pea performance across multiple physiological scales from nutrient acquisition to canopy chlorophyll status and photosystem function, leading to reduced biomass accumulation. Inoculation with *B. subtilis* effectively alleviated these negative impacts, restoring nutrient uptake, improving photosynthetic efficiency, and enhancing biomass accumulation. The underlying resilience mechanism appears to be multifactorial, encompassing direct nutrient mobilization, systemic effects, and rhizosphere microbiome restructuring, suggesting *B. subtilis* as a holistic modulator of pea performance under high pH conditions.

### *B. subtilis* rebalances physiological responses under alkaline stress

The present study highlights substantial genotype-dependent variation in alkaline stress tolerance and responsiveness to *B. subtilis* (Table 1). Although all cultivars experienced reduced growth and Chl content under alkaline conditions, the magnitude of recovery following inoculation differed markedly across genotypes. Such variability is consistent with previous reports, suggesting host genotype as a major player in shaping rhizomicrobial colonization, signal perception, and downstream transcriptional reprogramming (Thapa et al., 2025a; Jacoby et al., 2021). Genotype-specific differences in host response modulation and symbiotic compatibility likely influence the extent to which *B. subtilis* helps pea plants cope with stress. Alkaline stress reduced Fe and Mn concentrations in both roots and leaves in Sugar Snap, aligning with the reduced solubility and impaired Strategy I Fe acquisition in high-pH soils (Kim and Guerinot, 2007). In this study, *B. subtilis* restored Fe and Mn availability to near control levels and improved Ca and N accumulation, indicating coordinated recovery of nutrient homeostasis in peas. These ionomic recoveries paralleled gains in plant biomass and photosynthetic parameters, pointing to the coordinated restoration of resource supply and photochemical capacity. Such recovery aligns with established traits of *B. subtilis* relevant to alkaline soils, including siderophore and organic acid production, and antioxidant pathway activation (Kloepper et al., 2004; Egamberdieva et al., 2017). The fact that similar benefits occurred across genetically diverse pea cultivars, albeit with variable magnitudes, reflects the common but genotype-dependent nature of PGPR–legume interactions (Hasan et al., 2025; Vejan et al., 2016). Overall, these findings demonstrate that *B. subtilis* is highly effective in restoring micronutrient balance and photosynthetic performance in legumes under alkaline stress.

### Split-root responses are consistent with plant-wide effects

The split-root assay showed that *B. subtilis* inoculation of only part of the root system improved plant performance beyond the inoculated root compartment, indicating that its beneficial effects were not restricted to the local rhizosphere. These responses are consistent with systemic communication between root compartments; however, the signaling molecules involved were not directly investigated in this study. Previous studies have shown that microbe-induced plant-wide responses can involve mobile phytohormones, peptides, metabolites, and other vascular signals that coordinate physiological responses between spatially separated tissues (Martínez-Medina et al., 2013). Thus, the improved biomass and photosynthetic performance observed following localized inoculation are suggestive of plant-wide regulation rather than direct evidence of a specific systemic signaling pathway. Identification of the signals responsible for these responses will require targeted molecular and metabolomic analyses.

Such long-distance communication resembles induced systemic tolerance reported for other plant PGPR, where signal perception in one root region primes the rest of the plant for improved stress resilience (Conrath et al., 2006; Hartmann et al., 2021). Additionally, microbe-triggered systemic reprogramming often enhances antioxidant defenses, nutrient mobilization pathways, and the expression of transporter families critical for nutrient economy under abiotic stress (Verma et al., 2024; Bhattacharyya & Jha, 2012). Our transcriptomic enrichment in transporter and signaling genes aligned with this established framework, supporting the idea that *B. subtilis* activates a coordinated, plant-wide regulatory network rather than exerting effects solely on the site of inoculation. This systemic coordination also mirrors recent findings where PGPR elicit cross-compartmental benefits through volatile compounds, secreted metabolites, and immune-regulatory cues that act across the root–shoot continuum (Ryu et al., 2004; Fincheira & Quiroz, 2018). Together, our observations are consistent with a plant-wide response potentially involving long-distance signaling, although the underlying signals remain to be identified.

### Inorganic Fe reduces symptoms, but *B. subtilis* drives symbiotic recovery

The FeEDDHA supplementation study offered mechanistic control by directly supplying a highly stable Fe-chelate capable of maintaining Fe solubility at high pH (Rojas et al., 2008). Alkaline soils can impair root growth, thereby reducing the efficient absorption of essential minerals by plants (Saleem et al., 2023). In this study, while FeEDDHA remains stable under alkaline conditions, a portion of the Fe can still bind to soil particles, decreasing its bioavailability and overall effectiveness (Zuluaga et al., 2023). Although inorganic Fe restored chlorophyll content and partially improved plant growth, it did not recover nodulation or the rhizosphere Fe-chelating activity observed in *B. subtilis*-treated plants, highlighting its narrower mode of action compared with microbial inoculation. Once Fe limitation was chemically alleviated, the physiological requirement for *B. subtilis*-mediated nutrient mobilization was reduced. Nevertheless, inorganic Fe supplementation provided only modest improvements and did not fully overcome the constraints imposed by high pH. In particular, inorganic Fe alone was unable to restore nodulation or reproduce the enhanced Fe-chelating activity associated with *B. subtilis* treatment. Given that the CAS assay measures general Fe-chelating activity and does not identify specific siderophore compounds, these results should be interpreted as siderophore-associated rather than direct evidence of specific siderophore production. The consistently higher Fe-chelating activity in *B. subtilis*-treated plants suggests that this microbe may enhance Fe availability and rhizosphere function under alkaline stress through mechanisms that warrant further investigation. Collectively, these findings indicate that *B. subtilis* exerts broader beneficial effects than inorganic Fe amendment alone.

Since alkaline stress substantially inhibits nodulation, we next examined whether *B. subtilis* promotes *Rhizobium* performance directly, using co-culture assays. The co-culture experiments on Fe-sufficient plates revealed no significant differences in colony expansion between *B. subtilis* and *R. leguminosarum*. Under alkaline conditions, *R. leguminosarum* and *B. subtilis* respond differently to pH stress, resulting in distinct growth dynamics even within the same co-culture treatment. Such divergence did not appear under control conditions, indicating that the variation is possibly driven by species-specific sensitivity to alkaline media. However, co-culture at high pH enhanced *R. leguminosarum* growth, indicating a potentially favorable interaction between the two bacteria under alkaline conditions. A limitation of this study is that the co-culture assay was conducted on nutrient agar under controlled laboratory conditions, which may not fully capture the complexity of rhizosphere interactions occurring in soil. Microbial behavior and plant–microbe interactions in the rhizosphere are influenced by numerous biotic and abiotic factors that are not represented in agar-based systems (Mendes et al., 2013). Therefore, the co-culture results should be interpreted as evidence of potential microbial compatibility and interaction rather than a direct representation of rhizosphere dynamics. Since this interaction was demonstrated only *in vitro*, its contribution to nodulation and rhizobial performance in planta remains to be established. Also, future studies using soil-based and in planta systems will be necessary to further validate these interactions under more ecologically relevant conditions.

Consistent with this, improved Fe and N status supported by increased ammonium transport in *B. subtilis*-treated roots under alkaline stress suggests that these benefits are alkaline-dependent. Moreover, potential complementarity between *B. subtilis* and *R. leguminosarum* may contribute to improved nodulation and plant performance under alkaline conditions, although direct effects on rhizobial abundance and nitrogen fixation remain to be validated. This plant-mediated facilitation aligns with previous reports in PGPR–rhizobia systems, where enhanced symbiotic performance under stress relies more on the host’s metabolic and signaling reprogramming rather than on direct microbe–microbe interactions (Frey-Klett et al., 2011; Vishwakarma et al., 2020). This whole-plant response is consistent with transcriptomic enrichment in transporter and signaling genes that likely mediate these distal effects. These findings suggest a complementary role for microbial inoculants and inorganic Fe supplementation. FeEDDHA offers rapid correction of acute Fe deficiency, while *B. subtilis* provides a sustainable, broader-spectrum improvement that also benefits Mn and Ca uptake, photosynthetic resilience, and microbiome health. In low-input or organic systems, *B. subtilis* may thus serve as a viable alternative to synthetic chelates, avoiding their higher costs and potential environmental persistence. However, a limitation of this study is that alkaline stress was induced using a bicarbonate-based model, which captures key features of alkaline soils, including elevated pH and bicarbonate-associated nutrient constraints, but may not fully represent the physicochemical complexity of naturally occurring calcareous soils. Natural alkaline soils differ in mineral composition, carbonate content, salinity, organic matter, and microbial diversity, all of which can influence plant responses and nutrient dynamics (Farooqi et al., 2023; Zhang et al., 2008). Therefore, while the current system provides a controlled platform for mechanistic investigations, future field-based studies will be necessary to validate the effectiveness of B. subtilis under naturally alkaline soil conditions.

### Transcriptomic reprogramming improves nutrient acquisition and redox homeostasis

RNA-seq analysis revealed that *B. subtilis* inoculation under alkaline stress triggered a broad transcriptional reprogramming in pea roots, with pronounced upregulation of genes associated with symbiotic association, mineral acquisition, phytohormone signaling, and redox regulation. Alkali-exposed plants inoculated with *B. subtilis* showed the upregulation of several sugar transport–related genes (*SWEET*, *GLUT)*. Upregulation of multiple sugar transporters indicates enhanced carbon allocation and exchange during symbiotic association with *B. subtilis*. Such adjustments likely support microbial metabolism while maintaining the plant’s energy balance (Li et al., 2025; Julius et al., 2017). Also, these transporters likely facilitate bidirectional sugar movement between the host and symbiotic partners mediated by *B. subtilis*, ensuring an adequate supply of carbon for microbial metabolism while maintaining host energy balance. Particularly, SWEET sugar transporters not only facilitate carbon allocation but also modulate nodule functioning and microbial colonization (Chen et al., 2023; Breia et al., 2021). Efficient sugar trafficking can support carbon allocation to nodules and rhizosphere microbes and thereby facilitate symbiotic associations (Lei et al., 2025; Pieterse et al., 2014; Zamioudis & Pieterse, 2012). Such reprogramming of carbohydrate transport pathways aligns with reports that PGPR like *B. subtilis* stimulate sugar-mediated nutrient exchange to sustain nitrogen fixation and plant resilience under high pH conditions (Berendsen et al., 2018; Kryvoruchko et al., 2018). This may be linked to the upregulated genes for ammonium transport (*PSAT0S1495G0040*), amino acid transport (*PSAT7G178640*), and Cys/Met metabolism (*PSAT6G052400*) that are associated with S and S assimilation, critical for stress adaptation (Hasan et al., 2025). Furthermore, enrichment of transporter-related transcripts, including those for Fe, Mn, nitrate, and phosphate uptake, suggests an enhanced capacity for nutrient mobilization and assimilation under conditions where availability is otherwise restricted by high pH (Lucena, 2006; Kobayashi & Nishizawa, 2012). Ion transport–related genes, including cation/H exchanger (*PSAT3G088320*) and ATPase (*PSAT7G218840*), were also elevated, likely contributing to pH homeostasis and nutrient uptake under alkaline stress (Wang et al., 2014). ABC transporters (*PSAT2G058680* and *PSAT3G053400*) and oxoglutarate-related metabolism (*PSAT6G045440*) may further support metabolite exchange and carbon–nitrogen balance, as previously reported for legumes under symbiotic and abiotic stress conditions (Khan et al., 2020).

Furthermore, there was a strong induction of genes involved in redox homeostasis and osmotic adjustment, suggesting that *B. subtilis* enhances antioxidant defense and water balance under alkaline stress. Similarly, the oxidative stress regulator suggests coordinated activation of stress-responsive transcriptional programs, in agreement with other findings that PGPR treatments can activate redox signaling networks in legumes (Al-Turki et al., 2023; Ahmad et al., 2022). Upregulation of oxidoreductase genes likely bolsters the plant’s oxidative defense, enabling protection of cellular structures and photosynthetic machinery from oxidative damage (Choudhury et al., 2017). Elevated ferritin-like protein transcripts (*PSAT6G040040*) indicate improved Fe storage and mitigation of Fe-induced oxidative bursts (Hasan et al., 2025). In parallel, there was a suppression of defense-associated transcripts, indicating a strategic reallocation of resources away from constitutive defense toward growth and nutrient acquisition, a hallmark of induced systemic tolerance (Conrath et al., 2006). This transcriptional profile is consistent with the plant-wide responses observed in the split-root assay, although the present data do not directly establish the identity or mechanism of long-distance signals. Such a coordinated molecular shift is consistent with the multifaceted benefits of *B. subtilis* inoculation in garden pea, where stress tolerance and symbiotic compatibility converge to enhance overall plant performance under alkaline stress conditions.

### Microbiome restructuring as an ecological layer of alkaline tolerance

The PCoA analysis of bacterial communities demonstrated that *B. subtilis* inoculation modulated rhizosphere assemblages under alkaline stress. Alkaline stress altered bacterial richness patterns, and *B. subtilis* modestly shifted richness profiles, although diversity metrics were not fully restored. The similarity of bacterial community structure in non-stress conditions with or without *B. subtilis* suggests that *B. subtilis* does not disrupt native bacterial networks in healthy soils but rather coexists within the established community (Hartman & Tringe, 2019). In contrast, alkaline stress induced a substantial shift in community composition, likely driven by high pH-mediated alterations in nutrient availability for stress-tolerant taxa (Rousk et al., 2010). Importantly, the partial restoration of the microbial profiles under alkaline stress toward the control cluster implies that *B. subtilis* inoculation rebalanced the microbial network under stress, potentially via root exudate modulation and production of metabolites that favor beneficial taxa (Gong et al., 2023). At the genus level, the enrichment of *Pseudomonas* and *Pseudorhizobium* influenced by *B. subtilis* under alkaline stress aligns with their documented plant growth–promoting traits, including production of phytohormones, phosphatases, and siderophores (Compant et al., 2019; Glick, 2014). *Pseudomonas* spp. are well-known for producing siderophores and phytohormones under alkaline conditions (Lemare et al., 2022; Sainz-Mejías et al., 2020). Furthermore, studies demonstrate that fluorescent pseudomonads secrete high-affinity siderophores (pyoverdines) that mobilize Fe and can be especially beneficial under alkaline or Fe-limiting soils (Cornelis and Matthijs, 2007; Kümmerli, 2022). Many strains solubilize inorganic phosphate via the PQQ-dependent gluconic-acid pathway and phosphatases, improving P uptake and root growth (Lugtenberg & Kamilova, 2009). *Pseudorhizobium* is a genus separated from Rhizobium by phylogenomics, with isolates from seawater, rocks, and polluted soils, adapted to harsh environments (Lassalle et al., 2021). Although its effects on crops are less documented, *Pseudorhizobium’s* contribution to plant performance likely stems from metal/metalloid transformation and stress-tolerant physiology (Lassalle et al., 2021; Andres et al., 2013). The enrichment of *Klebsiella* and *Brevundimonas* due to *B. subtilis* under non-stress conditions points toward enhanced functional redundancy in the rhizosphere, which could buffer against future environmental perturbations. Collectively, these results indicate that *B. subtilis* inoculation is associated with restructuring of the bacterial community under alkaline stress, although the functional consequences of these shifts require further validation.

ITS-based profiling showed that *B. subtilis* moderated alkaline-induced shifts in fungal assemblages, partially realigning community composition toward the control state. Alkaline stress reduced fungal richness and diversity, consistent with the suppression of pH-sensitive saprophytic and mutualistic fungi (Lauber et al., 2009). Although richness and diversity did not differ significantly, fungal community composition shifted toward that of control treatments with BS inoculation*. Pseudallescheria* (teleomorph of *Scedosporium*) is best supported as a saprotroph that degrades lignin and aromatics, processes that can indirectly enhance nutrient turnover in soils (Poirier et al., 2023). Also, *Pseudallescheria* detected in plant tissues may reflect transient environmental contamination rather than stable, beneficial endophyte status (Zhang et al., 2023). However, *Chaetomium* can act as a beneficial endophyte/biocontrol (Feng et al., 2023). It can also trigger induced systemic resistance since transcriptomic work in tomato shows JA/ET-linked immune priming after *C. globosum* treatment (Singh et al., 2021). Beyond defense, *C. globosum* mobilizes phosphorus via secreted phosphatases/phytases, increasing crop P uptake and yield in wheat and pearl millet (Tarafdar and Gharu, 2006). Hence, the enrichment of *Pseudallescheria* and *Chaetomium* by *B. subtilis* under alkaline soil may provide indirect benefits to pea plants by accelerating the decomposition and recycling of nutrients. Although these taxa were enriched in association with *B. subtilis*-mediated recovery under alkaline stress, the present sequencing data do not establish their direct functional contribution to stress tolerance. Their enrichment may also reflect changes in root physiology or rhizosphere conditions induced by *B. subtilis*. Isolation and functional validation of these taxa will be required to determine their causal roles in alkaline stress tolerance. Further, the use of three biological replicates for RNA-seq and microbiome analyses may limit statistical power, particularly for detecting subtle treatment effects. Therefore, mechanistic relationships represent a working hypothesis integrating our experimental observations with proposed biological interactions, rather than directly demonstrated causal pathways (Fig. 9).

**Fig. 9.**
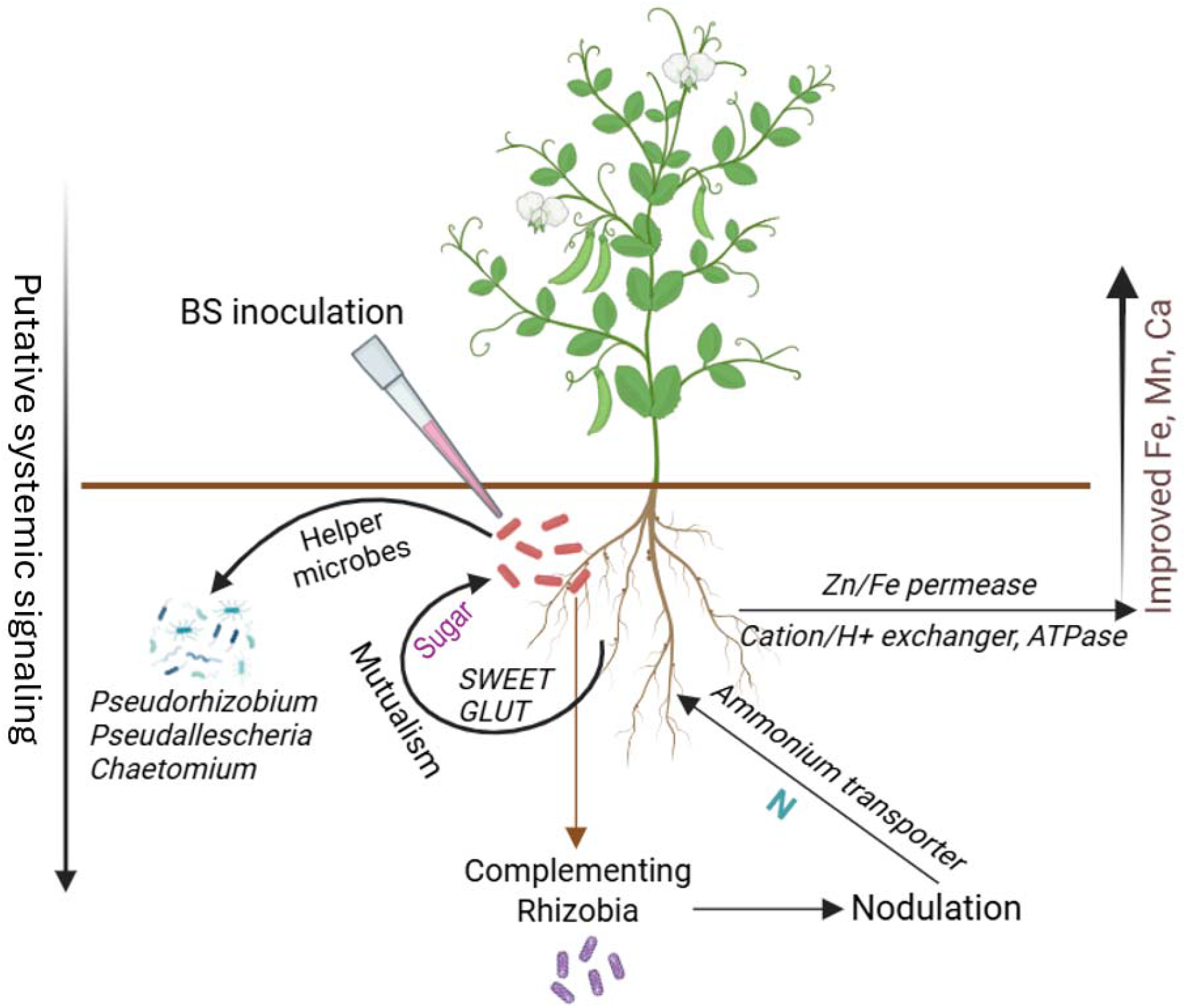
Hypothesized model of *B. subtilis*-associated responses contributing to alkaline stress tolerance in pea plants. *B. subtilis* inoculation was associated with enrichment of microbial taxa, including *Pseudorhizobium*, *Pseudallescheria*, and *Chaetomium*, alongside altered expressions of *SWEET* and *GLUT* transporters. These responses may contribute to changes in host–microbe interactions and nodulation, while increased expression of ammonium transporters and gene involved in nitrogen assimilation suggests altered N metabolism. In parallel, cation/H^+^ exchangers, ATPases, and Zn/Fe permeases may support nutrient homeostasis under alkaline stress. Together, these experimentally observed responses are consistent with, but do not directly demonstrate, putative systemic regulation contributing to improved nutrient status and plant performance under alkaline conditions.

## CONCLUSIONS

Our findings demonstrate that *B. subtilis* inoculation effectively mitigates the adverse effects of alkaline stress in garden pea by restoring Fe, Mn, and N levels, improving photosystem efficiency, and increasing biomass across diverse cultivars (Fig. 9). Split-root assays provided evidence for plant-wide responses consistent with systemic response, while co-culture experiments revealed growth-promoting interactions between *B. subtilis* and *Rhizobium*. The greater recovery of nodulation with *B. subtilis* than with inorganic Fe suggests that factors beyond Fe availability alone contribute to the observed symbiotic response. Transcriptome profiling further showed strong induction of genes associated with sugar transport, pH homeostasis, and nutrient assimilation, together with elevated expression of oxidoreductases and ferritin-like proteins, suggesting improved antioxidant defense and Fe storage following *B. subtilis* inoculation under alkaline stress. In parallel, *B. subtilis* enriched beneficial taxa, such as *Pseudomonas*, *Pseudorhizobium,* and *Chaetomium*, which are possibly enriched taxa to reinforce stress response under alkaline conditions. Collectively, these findings highlight how *B. subtilis* orchestrates molecular, physiological, and microbiome-level reprogramming, establishing its potential as a targeted bioinoculant for sustaining legume productivity in alkaline soils.

## Data availability

Illumina sequencing data were submitted to NCBI under the following accession numbers: PRJNA1313627 (RNA-seq), PRJNA1313634 (16S), and PRJNA1313644 (ITS).

## Acknowledgments

We express our gratitude to the Louisiana Biomedical Research Network for the funding. This research was also partially supported by a startup grant from Lamar University.

## Author contributions

AHK conceived the study, performed the molecular experiments, analyzed the data, performed bioinformatics analysis, and drafted the manuscript. AT cultivated the plants and performed a split-root assay. MRH performed microbial co-culture experiments and extracted the DNA. MGM interpreted the results and critically revised the manuscript.

